## Supplementary Information for "Competition contributes to both warm and cool range edges"

### Supplementary Methods

**Estimation of focal plant size.** To estimate focal plant size, we measured size-related morphological traits on all focal individuals at each census between 2017 and 2020 (i.e., the number and/or length of flowering stalks, leaves or ramets, depending on species) and predicted their dry aboveground biomass using regression models (mean $R^2$ = 0.871; Supplementary Data 1). These regression models were based on additional plants on which we measured morphological traits, and which were then oven-dried and weighed for dry biomass. We fitted seedling regression models using plants raised in the greenhouse to predict the size of plants which were initially transplanted into the field; adult plant models were fitted using plants taken from the background plots to estimate the size of non-flowering or flowering plants after at least one growing season. All models were fitted as zero-intercept models to avoid predicting negative plant sizes, except for *Crepis biennis*, for which non-zero intercept models were judged visually to provide better fits to the data.

**Estimation of focal plant seed production.** We fitted regression models between seed number and fruit length to predict seed number of focal plants. For each species, we measured ca. 20 × 3 sites = 60 intact flowers or fruits before seed-release at the early fruiting stage on background plants. Flowers and fruits were then air-dried, and seeds were counted. We first made a full model with site and its interaction with fruit length, which was further simplified to a species-level model by pooling the three sites together if the interaction was not significant. We fitted zero-intercept models to avoid negative estimates of seed number. We used the average number of seeds per fruit for species for which the number of seeds per fruit was observed to be invariant (*Salvia pratensis*) or for which seed number was poorly correlated with fruit size (*Anthylis vulneraria* ssp. *alpestris*, *Crepis biennis*, *Medicago lupilina*, *Trifolium badium*; Supplementary Data 2).

**Estimation of germination and recruitment.** To estimate germination and recruitment of each species, we conducted a separate experiment between 2018 and 2019. Within each site, we sowed seeds of 13 species (all except *Daucus carota*) in a 1 x 1 m plot filled with sterilized soil in autumn 2018. Seeds were glued to wooden toothpicks using water soluble glue and inserted ca.2 cm deep into the soil at 5 cm spacing. Each species had 20 sticks within each site resulting in ca. 240-1660 seeds per species depending on seed size. In the following spring, we counted germinated seedlings and then thinned them to leave a single individual per stick and a maximum of 10 sticks per species to avoid competition among plants. We followed those individuals until the end of the growing season to estimate seedling establishment. Note, we only measured recruitment in the absence of competition with this experiment. Since competition can substantially impact establishment (e.g., ref [1]), we approximated a competition-dependent recruitment rate for each species combination by multiplying the competition-free recruitment rates by survival rates of focal plants in competition plots; we restricted this analysis to focal plants that had been transplanted in spring and censused in their first growing season, to align these data with the germination experiment. We followed the same protocol to estimate germination and recruitment of *D. carota* between 2019 and 2020.

**Population modelling**. Integral projection models (IPM) model $\mathrm{n}(z1, t+1)$, the size distribution of the population at time $t+1$ as a function of $\mathrm{n}(z,t)$, the size distribution at time $t$ and $\mathrm{K}(z1,z)$, a kernel describing size-specific transitions from size $z$ to $z1$ through growth $\mathrm{P}(z1,z)$ and reproduction $\mathrm{F}(z1,z)$ [2, 3]:

$$\mathrm{n}(z1, t+1) = \int_L^U \mathrm{K}(z1,z)\mathrm{n}(z,t)\mathrm{d}z \quad (1)$$

$$= \int_L^U [\mathrm{P}(z1,z) + \mathrm{F}(z1,z)]\mathrm{n}(z,t)\mathrm{d}z \quad (2)$$

where *L* and *U* denote the lower and upper size bounds covering all possible sizes within the population. Here, we used a pre-reproduction census IPM for our perennial plants [3] and plant size (i.e., estimated dry aboveground biomass; log scale) as the continuous state variable, $z$. The growth kernel consists of survival and growth: $\mathrm{P}(z1,z) = \mathrm{s}(z)\mathrm{G}(z1,z)$, where $\mathrm{s}(z)$ is the probability of a plant of size $z$ surviving until the following year and $\mathrm{G}(z1,z)$ is the probability of a plant growing from size $z$ to $z1$. Fecundity is described by $\mathrm{F}(z1,z) = \mathrm{f}(z)\mathrm{b}(z)\mathrm{p_g}\mathrm{p_r}\mathrm{C1}(z1)$, where $\mathrm{f}(z)$ is flowering probability while $\mathrm{b}(z)$ is the number of seeds produced by a reproductive individual of size $z$. $\mathrm{p_g}$ and $\mathrm{p_r}$ are constant probabilities of seed germination and seedling recruitment, respectively, and $\mathrm{C1}(z1)$ is the offspring size distribution at time $t+1$ (Supplementary Table 3).

To investigate the effects of plant size, competitor species and site (elevation) on vital rates, we first fitted a full model including all possible main effects and their interactions. The full models of size-dependent vital rates (i.e., survival, growth, flowering and fecundity) included three main effects (size $z$ at time *t*, experimental site and competitor species) and all two- and three-way interactions, while the full models of size-independent vital rates (i.e., germination and recruitment) included only site, competitor species and their interaction (Supplementary Data 3&4). For each vital rate of each species, we then compared all nested models of full models using the Akaike information criterion corrected for small samples (AICc, Supplementary Data 4). The benefit of this approach is that we can borrow strength across treatments in cases where full modes were outperformed by reduced models. Note that we had to use a simpler model in cases where the lowest-AICc model appeared to be overfitting. We then obtained vital rate parameters from the best-fit models (Supplementary Data 3).

Given the potential effects on vital rates of plant positions within plots (e.g., edge effects) and non-independence of repeated measures across years on the same individuals, we also fitted mixed-effects models with plant position and identity of individual as random effects. However, this resulted in convergence and singularity issues for most vital rates models. In addition, we could not fit year as a random effect because it was confounded with plant size and age, given that we transplanted most focal seedling plants at the same time, though we acknowledge the potentially important effects of interannual variation in climate on demographic performance. Therefore, we dropped random effects for all vital rates and fitted IPMs using parameters obtained from fixed-effects models.

We estimated the mean and standard deviation of offspring size using seedlings raised in the greenhouse. We set the lower size limit $L$ as the minimum size of greenhouse seedlings corresponding to the size of plants that initiated the experiment. To determine the upper size limit, we first examined how size $z1$ was affected by site and competitor species by comparing models with size $z1$ as the response variable following the same AICc-based model selection as for vital rates. The upper size limit was then set as the maximum plant size observed in a given competitor species, site or both, depending on which terms were retained in the best-fitting model (Supplementary Data 3). Following the methods suggested by Ellner et al. [3], we found that the size limits determined in this way caused only a low level of size eviction (on average across all IPM models, 3% of individuals become evicted). We determined the number of bins by projecting IPMs starting with 100 bins and increased it until the projected population growth rates stabilized [3]. The projected population growth rates of most of IPMs stabilized with 3000 bins, which was then used for subsequent analysis.

**Flowering phenology and overlap.** The flowering of background plants (the percent of plants that had flower buds or open flowers) was recorded weekly during the growing season of 2019 (April to August) for each species at each site. To determine the date of the first and last flowering of each species in each site, we fitted generalized additive models (GAMs) using the relative abundance of flowering plants varying with the day of the year (Supplementary Fig. 7). For a given pair of species (same pairs as in the competition experiment), the temporal overlap of flowering phenology was then calculated as the number of days when both species flowered divided by the total length of the flowering period (i.e., the number of days between the earliest and latest flowering of the two species following ref. [4]).

**Supplementary Table 1**. Environmental characteristics of the three study sites. Long-term mean annual temperature was derived from climate interpolations based on data from 1981-2015 [5]. Mean annual temperature of air (15 cm above ground surface) and soil (8 cm below ground surface), and mean soil water content (8 cm below ground surface) were recorded using TMS-4 dataloggers (TOMST®, Czech Republic; https://tomst.com/web/en/) and averaged between July 2019 to July 2020. The number of days covered by snow were calculated as the number of days of which daily mean temperature at the ground surface was between -0.5 and 0.5 °C.

| **Variable** | **Les Posses** | **Solalex** | **Anzeindaz** |
|---|---|---|---|
| Elevation (m a.s.l.) | 890 | 1400 | 1900 |
| Latitude (°N) | 46.2706 | 46.2866 | 46.2875 |
| Longitude (°E) | 7.0327 | 7.1338 | 7.1687 |
| Long-term mean annual temperature (°C) | 9.6 | 5.9 | 2.5 |
| Mean annual air temperature (°C) | 11.4 | 7.9 | 5.9 |
| Mean annual soil temperature (°C) | 10.4 | 6.5 | 4.9 |
| Mean volumetric soil water content | 0.261 | 0.410 | 0.438 |
| Number of days under snow | 12 | 129 | 163 |

**Supplementary Table 2.** Species included in this study. The elevation range is defined as the 10th and 90th percentile of a species' elevation distribution in the study area (see Methods). *Anthyllis vulneraria* has a broad range, but the alpine ssp. *alpestris* (which was not differentiated in the available distribution data [6]) was used for the experiments.

| Species | Code | Family | Functional group | Origin of elevation | Growth form | Elevation range (m, mean, lower-upper) | Seed supplier |
|---|---|---|---|---|---|---|---|
| Bromus erectus | Brer | Poaceae | Grass | Lowland | Perennial | 971, 598-1351 | Otto Hauenstein Samen |
| Crepis biennis | Crbi | Asteraceae | Forb | Lowland | Biennial | 1007, 764-1299 | Otto Hauenstein Samen |
| Daucus carota | Daca | Apiaceae | Forb | Lowland | Biennial | 1063, 683-1429 | UFA SAMEN |
| Medicago lupulina | Melu | Fabaceae | Legume | Lowland | Perennial | 1035, 653-1408 | Otto Hauenstein Samen |
| Plantago lanceolata | Plla | Plantaginaceae | Forb | Lowland | Perennial | 1169, 629-1657 | UFA SAMEN |
| Poa trivialis | Potr | Poaceae | Grass | Lowland | Perennial | 980, 527-1390 | UFA SAMEN |
| Salvia pratensis | Sapr | Lamiaceae | Forb | Lowland | Perennial | 782, 539-1069 | Otto Hauenstein Samen |
| Anthylis vulneraria ssp. alpestris | Anal | Fabaceae | Legume | Highland | Perennial | 1848, 1341-2217 | Kaertner Saatbau |
| Arnica montana | Armo | Asteraceae | Forb | Highland | Perennial | 1906, 1622-2091 | Jellito |
| Aster alpinus | Asal | Asteraceae | Forb | Highland | Perennial | 2108, 2002-2236 | Jellito |
| Plantago alpina | Plal | Plantaginaceae | Forb | Highland | Perennial | 1916, 1581-2193 | Schutz Filisur |
| Poa alpina | Poal | Poaceae | Grass | Highland | Perennial | 2045, 1674-2458 | Kaertner Saatbau |
| Sesleria caerulea | Seca | Poaceae | Grass | Highland | Perennial | 2013, 1652-2371 | Jellito |
| Trifolium badium | Trba | Fabaceae | Legume | Highland | Perennial | 1943, 1640-2253 | Schutz Filisur |

**Supplementary Table 3**. Structure and parameters of integral projection models (IPMs).

| Vital rate | Variable type | Distribution | Model | Meaning |
|---|---|---|---|---|
| Survival | Binary | Binomial | $\text{logit}[s(z)] = s_0 + s_1 * z + \sigma_s$ | Survival probability of plant of size z (log scale); $s_0$, $s_1$ and $\sigma_s$ are intercept, slope, and standard deviation of the linear function. |
| Growth | Continuous | Gaussian | $G(z1, z) = \frac{1}{\sqrt{2\pi\sigma_g^2}} \exp\left(-\frac{[z1 - \mu(z)]^2}{2\sigma_g^2}\right)$ Where $\mu(z) = g_0 + g_1 * z + \sigma_g$ | Probability of a plant growing from size $z$ to size $z1$ (log scale); $g_0$, $g_1$ and $\sigma_g$ are intercept, slope, and standard deviation of the linear function; $\mu(z)$ is expected size in year $t+1$ of a plant of size $z1$ in year $t$. |
| Flowering | Binary | Binomial | $\text{logit}[f(z)] = f_0 + f_1 * z + \sigma_f$ | Flowering probability of plant of size $z$ (log scale); $f_0$, $f_1$ and $\sigma_f$ are intercept, slope, and standard deviation of the logistic function. |
| Seed production | Integer | Gaussian | $b(z) = b_0 + b_1 * z + \sigma_b$ | Number of seeds produced by a plant of size z (log scale); $b_0$, $b_1$ and $\sigma_b$ are intercept, slope, and standard deviation of the linear function. |
| Germination | Continuous | Constant | $P_g$ | Constant probability of seed germination. |
| Seedling establishment | Binary | Binomial | $\text{logit}(P_r) = r_0 + \sigma_e$ | $r_0$ and $\sigma_e$ are intercept and standard deviation of the intercept-only logistic function. |
| Offspring size distribution | Continuous | Gaussian | $C1(z1) = \frac{1}{\sqrt{2\pi\sigma_r^2}} \exp\left(-\frac{(z1 - \mu_r)^2}{2\sigma_r^2}\right)$ | Probability of an offspring of size $z1$ (log scale); $\mu_r$ and $\sigma_r$ are mean and standard deviation of offspring sizes. |

**Supplementary Table 4.** Statistical analysis of intrinsic and invasion population growth rates using mixed-effect models (see Methods). *F* and *P* values were derived from likelihood ratio tests for the effects of elevation of the three sites (continuous), origin of focal species (lowland or highland) and their interactions on intrinsic and invasion growth rates based on the mean of 500 bootstrap replicates of the dataset. We included focal species as random effects ($n$ = 14). The total number of observations ($n$) is also shown. Significant terms are marked in bold.

| | | **Intrinsic population growth** | | **Invasion population growth** | |
|---|---|---|---|---|---|
| Source | d.f. | *F* | *P* | *F* | *P* |
| Elevation | 1 | 0.247 | 0.619 | 0.024 | 0.878 |
| Origin focal | 1 | **3.798** | **0.051 (.)** | **3.825** | **0.051 (.)** |
| Elevation: origin focal | 1 | **7.062** | **0.008 **** | **21.215** | **4.106e-06 ***** |
| *n* | | 41 | | 300 | |

*** $P < 0.001$; ** $P < 0.01$; * $P < 0.05$; (.) $P < 0.1$.

**Supplementary Table 5.** Statistical analysis of the coexistence metric, niche differences, and relative fitness differences using mixed-effect models (see Methods). *F* and *P* values were derived from likelihood ratio tests for the effects of elevation of the three sites (continuous), competitor identity (lowland-lowland, lowland-highland, and highland-highland) and their interaction on the average of the 500 bootstrap replicates of the dataset. For relative fitness differences, we also performed separate tests for each of the three types of pairs. We included the identity of species pairs as a random effect in all models. Also shown are the total number of observations (*n*) and number of pairs in parentheses. Significant terms are marked as bold. NA indicates the factor was not relevant in the models.

| | | **Coexistence metrics** | | **Niche differences** | | **Relative fitness differences** | | | | | | | |
|---|---|---|---|---|---|---|---|---|---|---|---|---|---|
| | | | | | | Overall | | low-low | | low-high | | high-high | |
| Source | d.f. | *F* | *P* | *F* | *P* | *F* | *P* | *F* | *P* | *F* | *P* | *F* | *P* |
| Elevation | 1 | **4.312** | **0.038** * | 1.547 | 0.213 | 1.406 | 0.236 | 0.192 | 0.661 | **3.979** | **0.046** * | **5.179** | **0.023*** |
| Competitor identity | 2 | 2.056 | 0.358 | 1.888 | 0.214 | 0.339 | 0.844 | NA | NA | NA | NA | NA | NA |
| Elevation: Pair type | 2 | **11.011** | **0.004** ** | **11.603** | **0.003** ** | 2.935 | 0.231 | NA | NA | NA | NA | NA | NA |
| *n* (number of pairs) | | 90 (38) | | 90 (38) | | 90 (38) | | 29 (10) | | 44 (20) | | 17 (8) | |

*** *P* < 0.001; ** *P* < 0.01; * *P* < 0.05; (.) *P* < 0.1.

**Supplementary Figure 1**. Field competition experiment. We interacted 14 species originating from low and high elevations in three sites across an elevational gradient in the western Swiss Alps. Within each site, focal plants were transplanted into non-competition (i.e., bare soil; grey) and competition plots, that is the established monocultures of lowland (orange) and highland (blue) species. Each species interacted with eight heterospecific species (interspecific pairs), including four lowland and four highland species, and itself (intraspecific pairs).

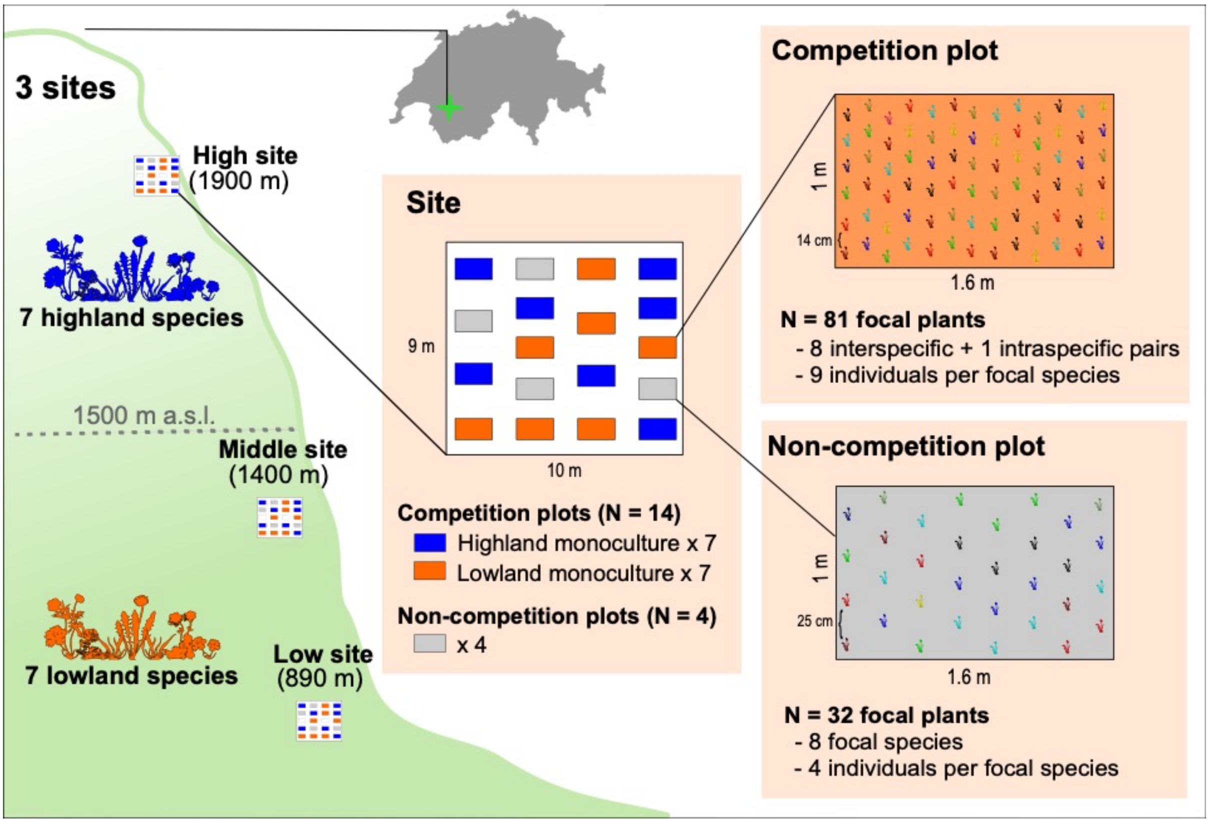

**Supplementary Figure 2.** Ambient temperature (**a**) and soil moisture (**b**) measured between July 2019 – October 2020 in the three study sites. Temperature of the air (15 cm above ground), ground surface and soil (8 cm below ground) and soil moisture (8 cm below ground) were recorded using TMS-4 dataloggers (TOMST®, Czech Republic; https://tomst.com/web/en/). Volumetric soil moisture was calibrated by using parameters of silt loam soil [7], which was used in the experiment.

**a**

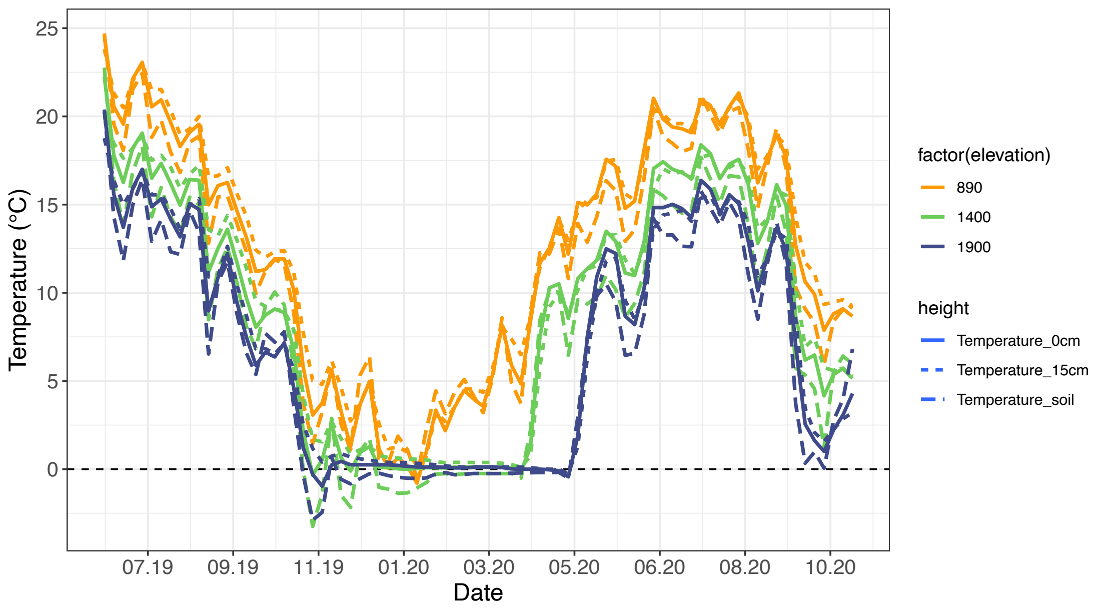


**b**

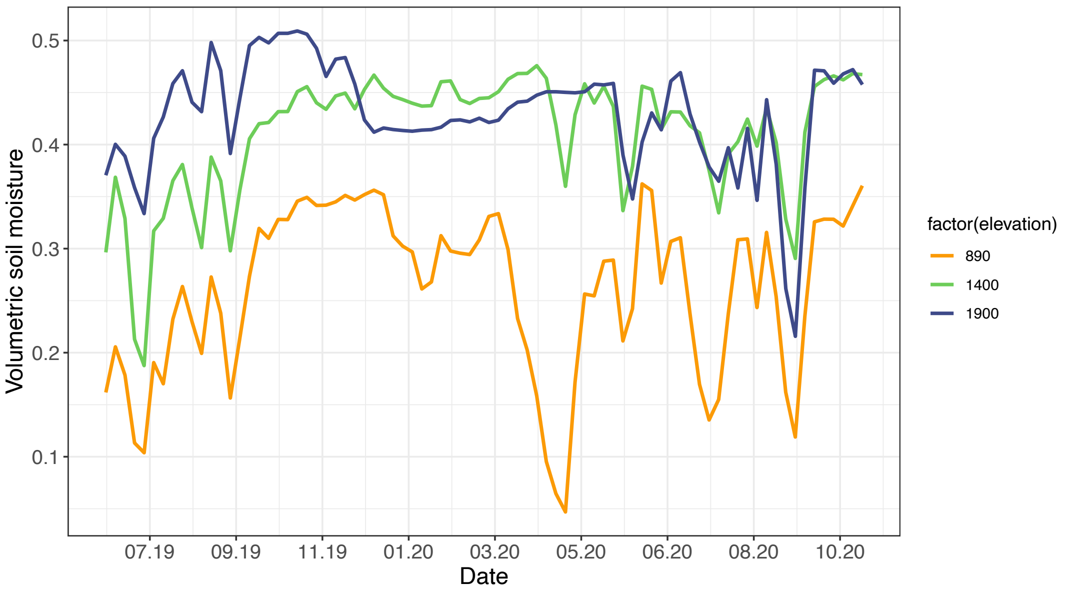

**Supplementary Figure 3**. Relationships between actual aboveground dry mass (y-axis) of training samples and estimated size (x-axis) using regression models. Each panel represents a single focal species. Point colors indicate different types of regression models used to predict size (see Supplementary Methods). Solid lines are the 1:1 line. See Supplementary Table 2 for species codes.

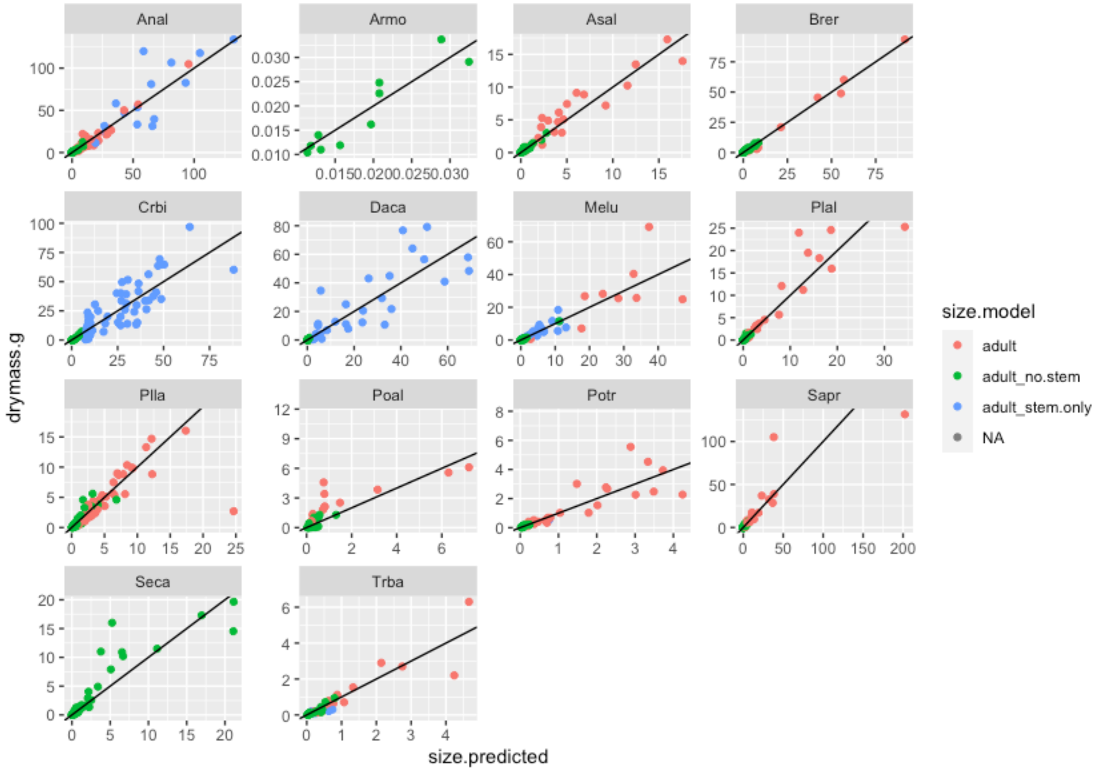

**Supplementary Figure 4**. The vital rates of each focal species growing with different competitors and at different sites. Vital rates are either size-dependent (**a**, survival; **b**, growth; **c**, flowering; **d**, fecundity) or size-independent (**e**, seed germination; **f**, seedling establishment in the absence of neighbours; **g**, seedling establishment under competition; **h**, recruit size distribution). For each vital rate, highland focal species are in the upper row and lowland species in the lower row. Grey points and bars in the background are observed values (see Supplementary Data 3 for the sample size of each vital rate and species), colored lines represent fitted vital rates implemented in the IPM models. Colours represent different competitor species (black for non-competition), line types represent sites (low, solid; middle, dotted; high, dashed). Overlapping lines and points indicate that the vital rates did not differ significantly between sites or competitor species based on model selection (See Supplementary Methods). The size range of the fitted lines represent the size bounds implemented in the IPM models (*L* and *U*; see Supplementary Methods). Rugs at the bottom indicate data distributions. Points were jittered for visual clarity for the probability of survival, flowering, and establishment. See Supplementary Table 2 for species codes.

**a**

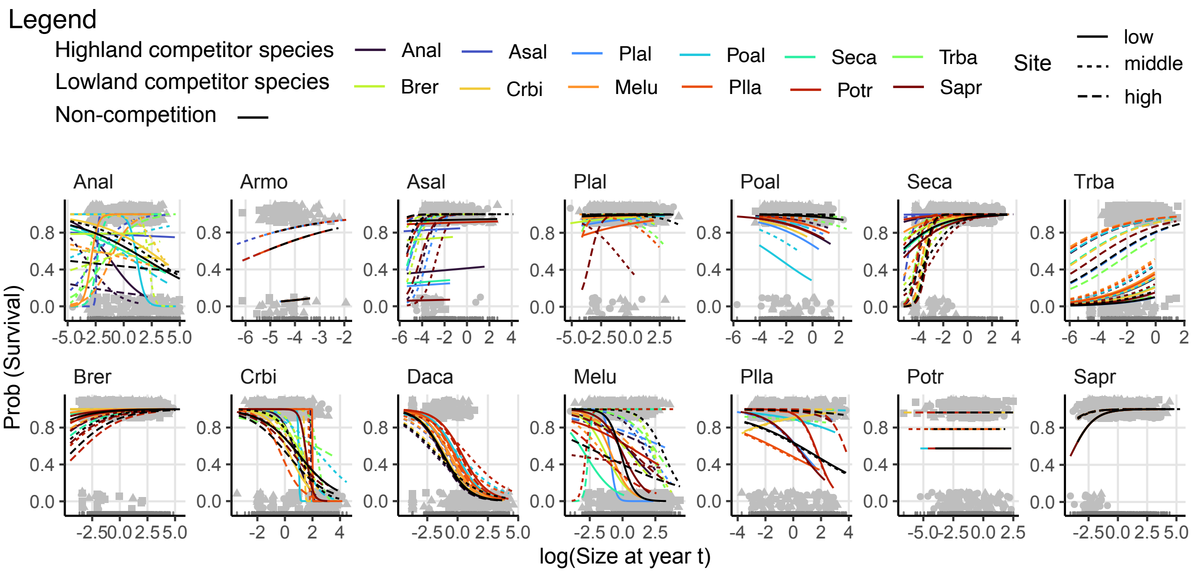


**b**

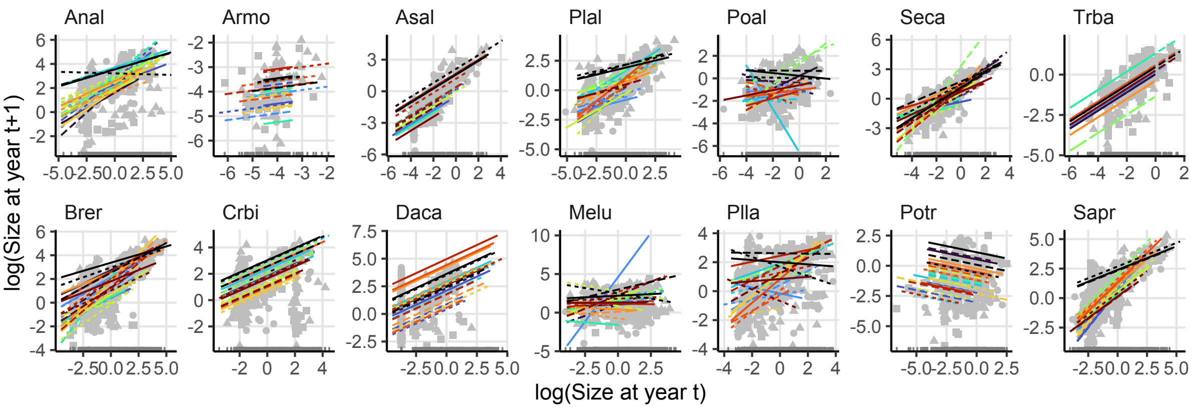

**c**

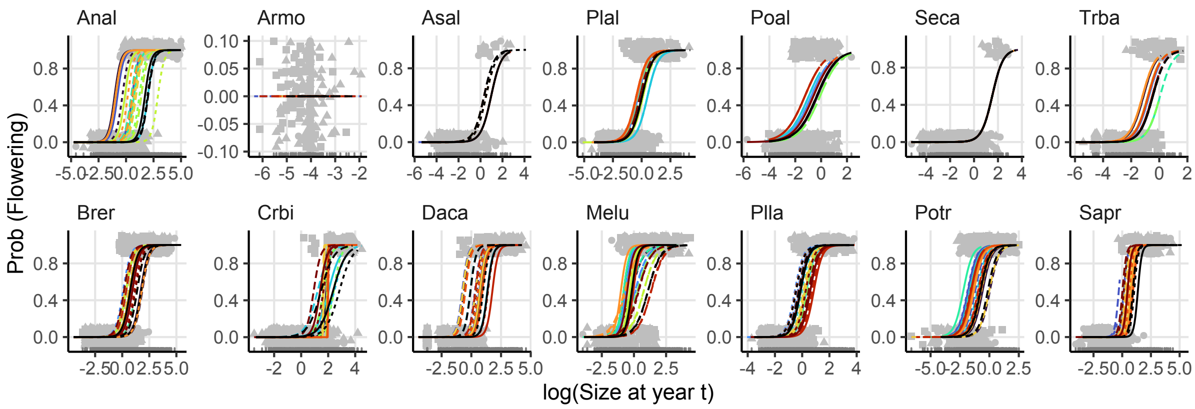
Anal
Armo
Asal
Plal
Poal
Seca
Trba
Brer
Crbi
Daca
Melu
Plla
Potr
Sapr
Prob (Flowering)
log(Size at year t)

**d**

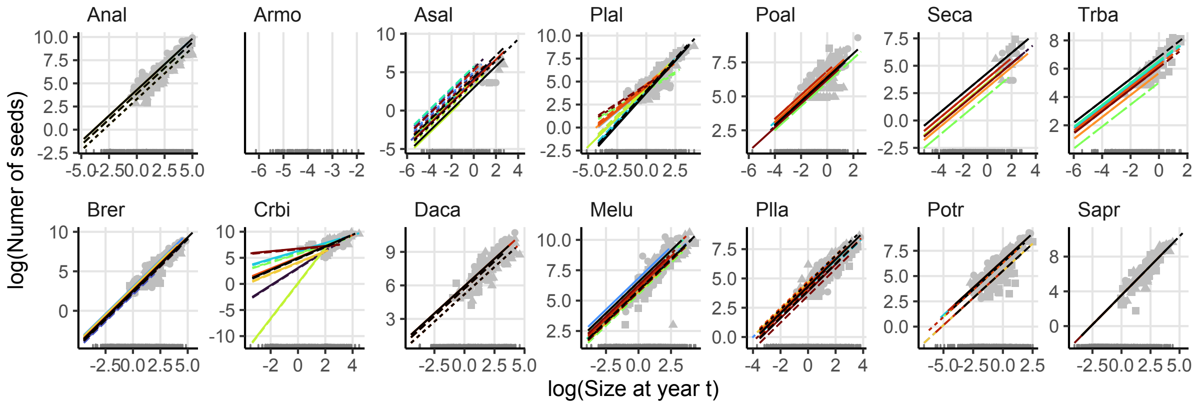
Anal
Armo
Asal
Plal
Poal
Seca
Trba
Brer
Crbi
Daca
Melu
Plla
Potr
Sapr
log(Numer of seeds)
log(Size at year t)

**e**

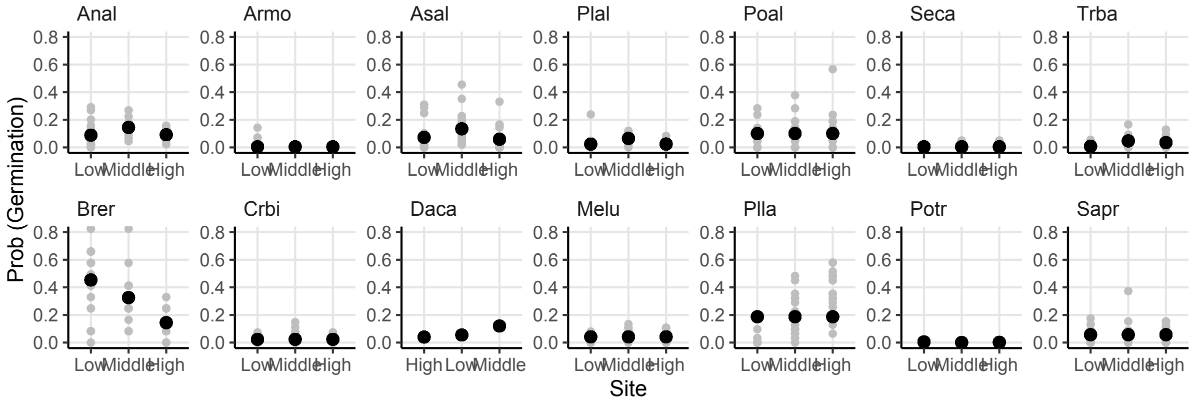
Anal
Armo
Asal
Plal
Poal
Seca
Trba
Brer
Crbi
Daca
Melu
Plla
Potr
Sapr
Prob (Germination)
Low
Middle
High
Site

**f**

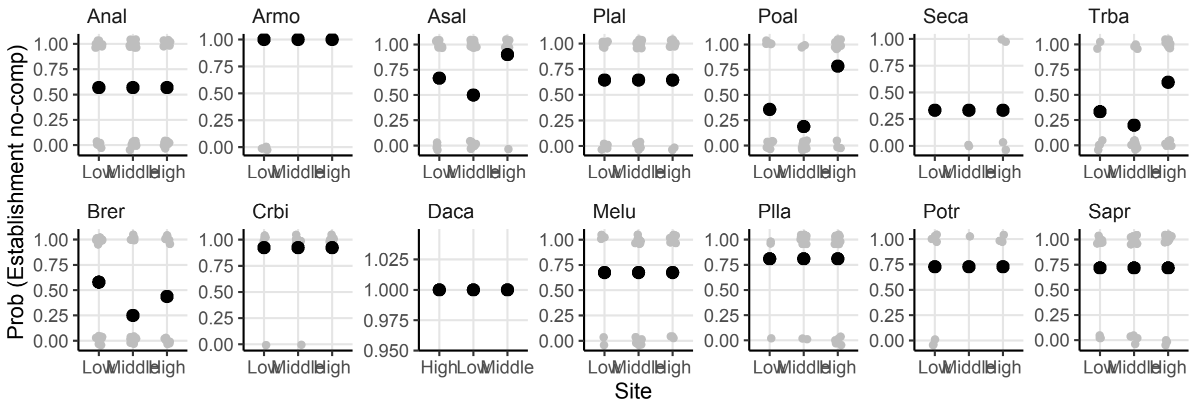
Prob (Establishment no-comp)
Anal
Armo
Asal
Plal
Poal
Seca
Trba
Brer
Crbi
Daca
Melu
Plla
Potr
Sapr
Site

**g**

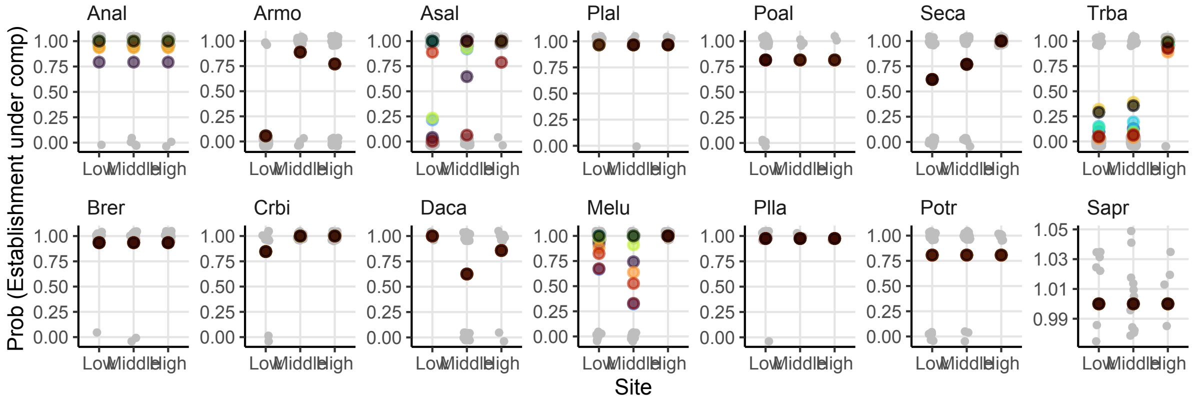
Prob (Establishment under comp)
Anal
Armo
Asal
Plal
Poal
Seca
Trba
Brer
Crbi
Daca
Melu
Plla
Potr
Sapr
Site

**h**

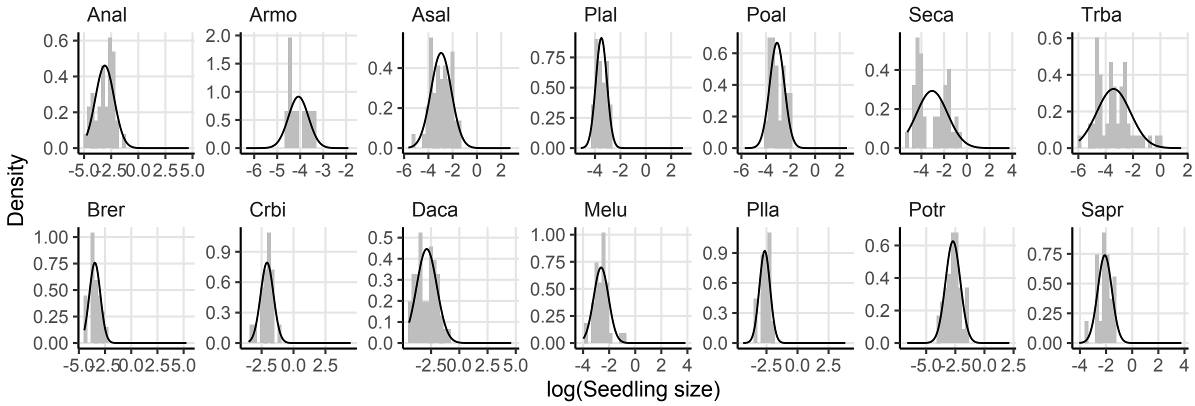
Density
Anal
Armo
Asal
Plal
Poal
Seca
Trba
Brer
Crbi
Daca
Melu
Plla
Potr
Sapr
log(Seedling size)

**Supplementary Figure 5**. Estimated population growth rates ($\lambda$) of lowland and highland species when invading conspecific monocultures (i.e., with intraspecific competition). The points and error bars represent the median and 95% confidence intervals based on 500 bootstrap replicates of the dataset. We expect that for populations (background monocultures) that are at their equilibrium abundance, ln($\lambda$) of conspecifics will be zero. The graphs show that the 95% confidence intervals for 32% (11 of 34) of intraspecific invasion growth rates (y-axis, log-transformed) included zero, implying that these monocultures were approximately at their equilibrium abundance. Ten monocultures were predicted to be above equilibrium abundance (ln($\lambda$) < 0) and 13 below equilibrium abundance (ln($\lambda$) > 0). See Supplementary Table 2 for species codes.

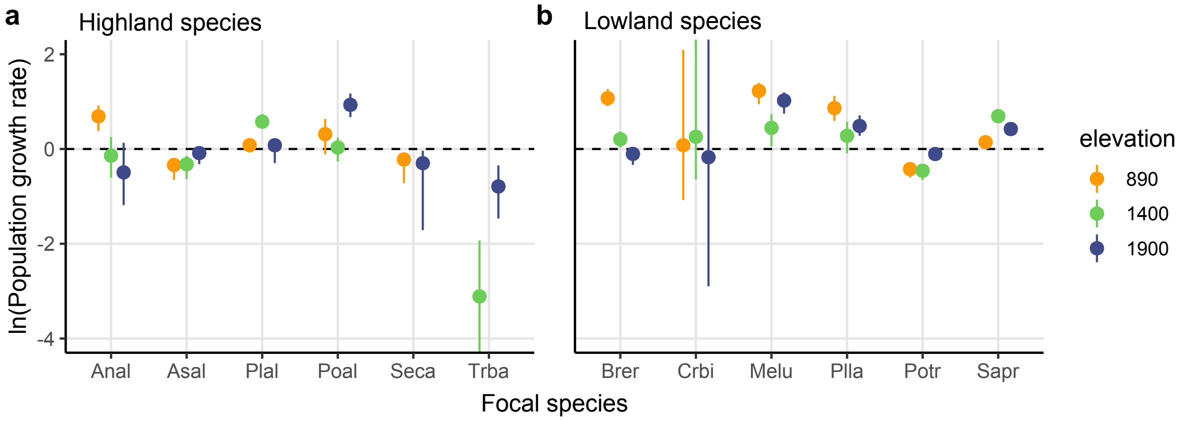

**Supplementary Figure 6**. Individual estimates of population growth rates ($\lambda$) for each focal species. Each panel is a single focal species growing at the three sites in the absence of competitors (i.e., intrinsic growth rates; green) and invading established monocultures of interspecific (black) and intraspecific (orange) competitors (i.e., invasion growth rates). Points and error bars represent the mean and 95% confidence intervals based on 500 bootstraps. Populations are predicted to grow when ln($\lambda$) > 0 (dotted line), and otherwise to decline. See Supplementary Table 2 for species codes.

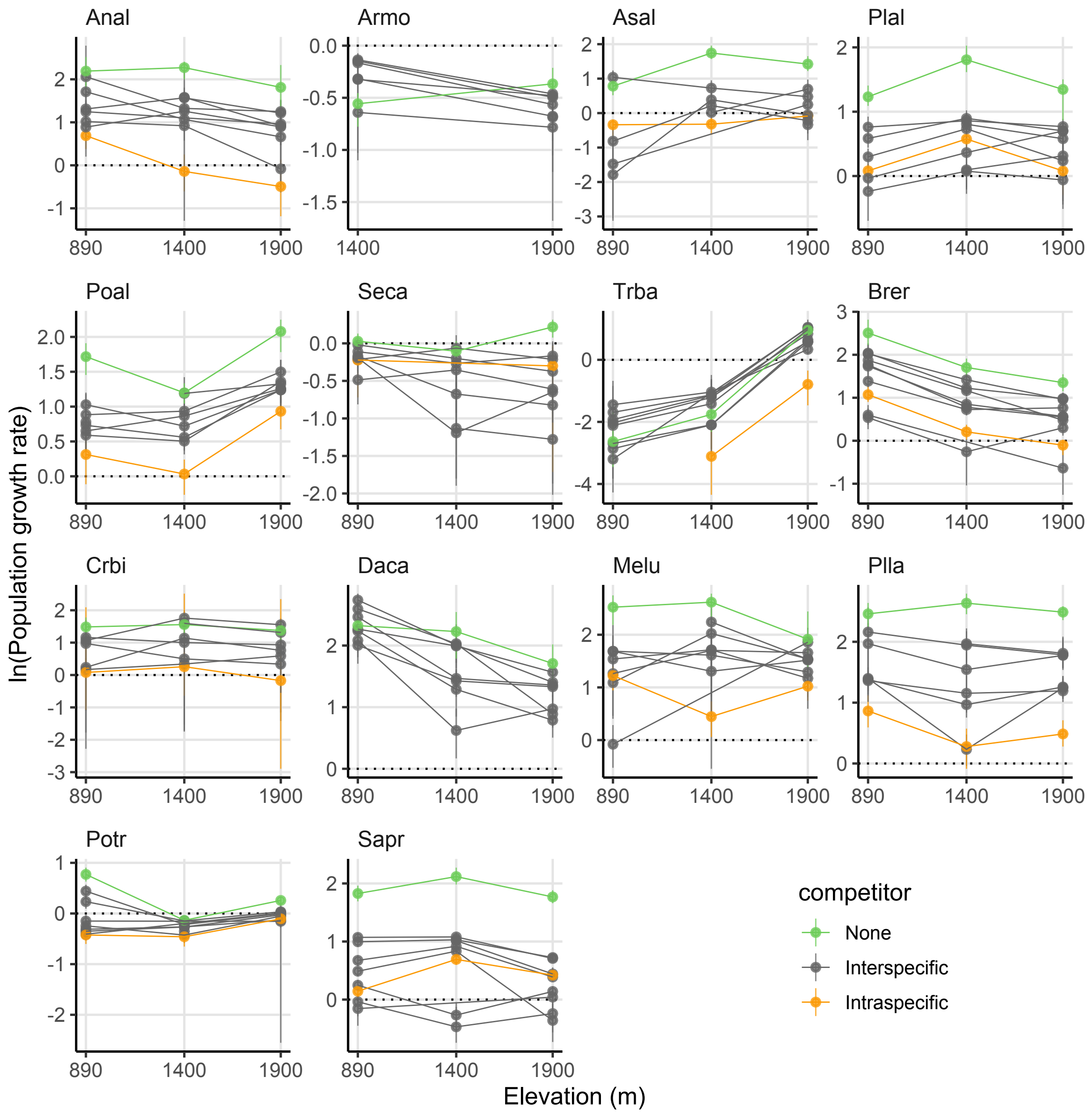

**Supplementary Figure 7**. Flowering phenology of lowland (orange tones, top row) and highland (green tones, bottom row) species in the low (left column), middle (middle column), and high (right column) sites. Phenological overlap of lowland or highland species at each site is shown on each panel. See Supplementary Table 2 for species codes.

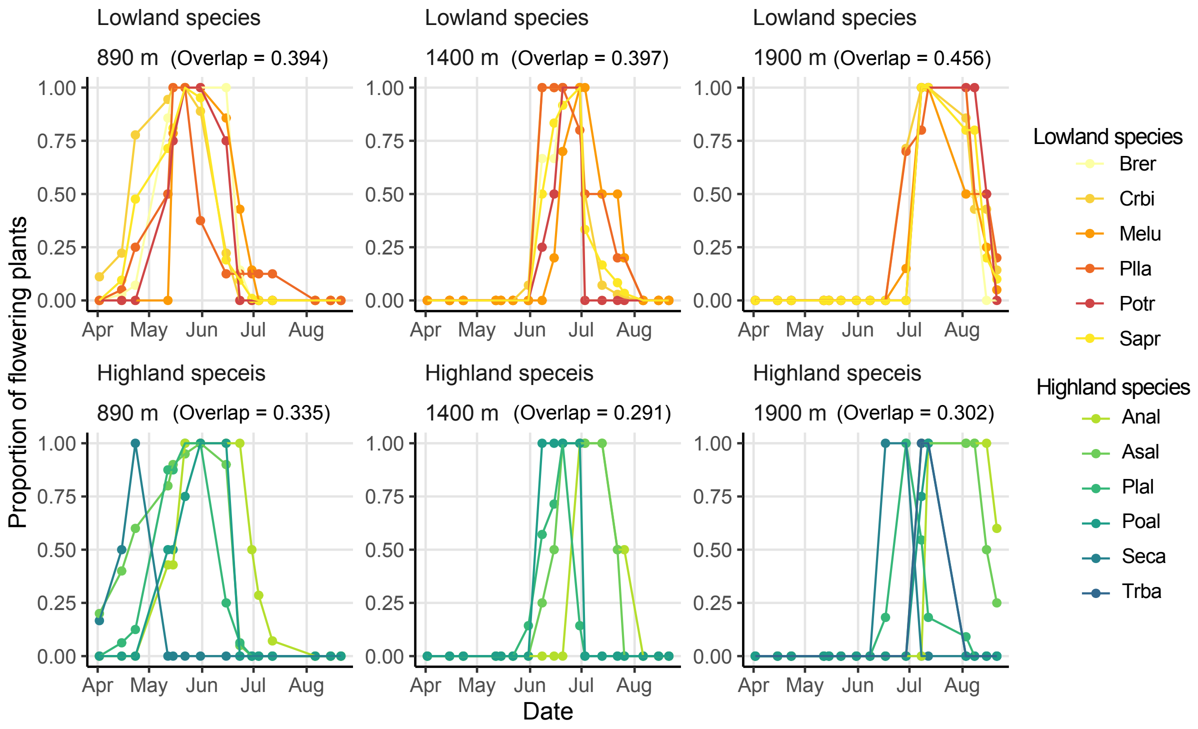
